# The importance and adaptive value of life history evolution for metapopulation dynamics

**DOI:** 10.1101/179234

**Authors:** Dries Bonte, Quinten Bafort

**Author notes:** 003291645213 - Corresponding author.

## Abstract

1. The spatial configuration and size of patches influence metapopulation dynamics by altering colonisation-extinction dynamics and local density-dependency. This spatial forcing as determined by the metapopulation typology then imposes strong selection pressures on life history traits, which will in turn feedback on the ecological metapopulation dynamics. Given the relevance of metapopulation persistence for biological conservation, and the potential rescuing role of evolution, a firm understanding of the relevance of these eco-evolutionary processes is essential.
2. We here follow a systems modelling approach to quantify the importance of spatial forcing and experimentally observed life history evolution for metapopulation demography as quantified by (meta)population size and variability. We therefore developed an individual based model matching an earlier experimental evolution with spider mites to perform virtual translocation and invasion experiments that would have been otherwise impossible to conduct.
3. We show that (1) metapopulation demography is more affected by spatial forcing than by life history evolution, but that life history evolution contributes substantially to changes in local and especially metapopulation-level population sizes, (2) extinction rates are minimised by evolution in classical metapopulations, and (3) evolution is optimising individual performance in metapopulations when considering the importance of more cryptic stress resistance evolution.
4. Ecological systems modelling opens up a promising avenue to quantify the importance of eco-evolutionary feedbacks for larger-scale population dynamics. Metapopulation sizes are especially impacted by evolution but its variability is mainly determined by the spatial forcing.
5. Eco-evolutionary dynamics can increase the persistence of classical metapopulations. The maintenance of evolutionary dynamics in spatially structured populations is thus not only essential in the face of environmental change; it also generates feedbacks that impact metapopulation persistence.

**Data-archive:** Metadata population dynamics in artificial metapopulations data are available from the Dryad Digital Repository: http://dx.doi.org/10.5061/dryad.18r5f (De Roissart, Wang & Bonte 2015). Modeling code is available on: https://github.ugent.be/pages/dbonte/eco_evo-metapop/ (ODD protocol in supplementary material)

## Introduction

Many, not to say all, species inhabit spatially heterogeneous and structured habitat. Spatial structure is an inherent property of natural systems because of spatial variation in the availability of diverse niche components, or because of human-induced fragmentation (Fahrig, 2003). When species move among patches of suitable habitat, local demography will be coupled with global metapopulation dynamics (Wang, Haegeman, & Loreau, 2015) and will thereby impact population size fluctuations and colonization-extinction dynamics (Cheptou, Hargreaves, Bonte, & Jacquemyn, 2017). Because the nature of these dynamics affects individual components of fitness, they are in turn anticipated to impose a strong selection on life history traits (Bonte & Dahirel, 2017; Olivieri, Couvet, & Gouyon, 1990).

To date, metapopulation viability is predominantly studied from an ecological perspective as human pressures impose changes in habitat, matrix composition, population sizes and connectedness (Bullock et al. 2018). These factors immediately impact the demographic properties of metapopulations (Hanski & Gilpin, 1997). Increased patch isolation does for instance increase dispersal mortality; local patch quality affects local growth rates; and patch size will eventually determine local carrying capacities and population sizes. High local population size, their summed metapopulation size and spatial asynchrony in local population fluctuations are generally anticipated to maximise metapopulation long-term survival and thus persistence (Hanski, 1998; Molofsky & Ferdy, 2005; Wang et al., 2015). The typology of the spatial habitat configuration will thus determine the demographic dynamics and eventually affect several components of metapopulation functioning. We refer to these ecological drivers as *spatial forcing*.

The spatial structure of metapopulations does equally impact trait evolution at the local and metapopulation-level (Cheptou et al., 2017). Such evolutionary consequences are mainly studied from a theoretical perspective with a strong focus on the evolution of dispersal and dispersal syndromes as these are per definition evolving at the larger metapopulation scale. The basic premise of these studies is that evolutionary changes in dispersal directly feedback on connectivity, hence enhancing metapopulation persistence (Bonte et al. 2018). Other traits have, however, been documented to evolve in parallel, and often in unpredictable ways (Bonte & Dahirel, 2017). Few empirical studies tested theoretical predictions beyond dispersal evolution, but recent work provides evidence of multivariate life-history evolution in metapopulations (De Roissart et al., 2016; Duplouy, Ikonen, & Hanski, 2013). Because often contrasting selection pressures act at the individual and metapopulation-level (Aspi, Jäkäläniemi, Tuomi, & Siikamäki, 2003; Laiolo et al., 2012), a maximised individual fitness will not necessarily lead to an optimised metapopulation persistence. It is for instance known that dispersal is not evolving to the best for the metapopulation, thus not evolving to such levels that performance in terms of metapopulation size or survival is maximised (Delgado, Ratikainen, & Kokko, 2011). In the extreme, evolution may induce evolutionary suicide, for instance by selecting against dispersal under extreme fragmentation, thereby compromising conservation efforts (Olivieri, Tonnabel, Ronce, & Mignot, 2016). Such scenarios occur for instance when habitat fragmentation selects against dispersal, thereby imposing a negative feedback between patch occupancy, inbreeding and extinction risk. An evolutionary perspective is thus highly relevant for contemporary biological conservation (Kinnison & Hairston, 2007).

Metapopulation persistence is affected by individual life history variation (Bonte & Dahirel, 2017). Because life history evolution takes place at the same temporal scale as changes in habitat fragmentation and the resulting demography (Cheptou et al., 2017), a tight feedback between the ecological and evolutionary dynamics (eco-evolutionary dynamics; [Hendry; intro this issue]; Bonte, Masier & Mortier 2018) is expected. The long-term research on the Glanville fritillary for instance demonstrated the tight eco-evolutionary coupling between dispersal evolution, local performance and colonisation dynamics to maintain metapopulation persistence (Fountain, Nieminen, Sirén, Wong, & Hanski, 2016; Hanski & Mononen, 2011; Hanski, Mononen, & Ovaskainen, 2011; Hanski & Saccheri, 2006). While we have some insights on the importance of spatial forcing for the evolution of specific traits, we lack any understanding on its feedback on metapopulation functioning, except for some theoretical work focusing on the eco-evolutionary feedback with range expansions and patch occupancy [Govaert et al. this special issue].

Several statistical and empirical approaches have been developed to quantify the relative contribution of ecology and evolution in demographic and phenotypic change. These approaches rely either on experimental manipulations of genetic variation and, hence, evolvability (Becks, Ellner, Jones, Hairston, & Hairston Nelson G., 2010; Turcotte, Reznick, & Daniel Hare, 2013; Turcotte, Reznick, & Hare, 2011; Van Petegem, Moerman, Dahirel *et al.* 2018), or on the statistical partitioning of time series data based on price and breeder equations or the use of Integral Projection Matrices (Coulson, 2012; Coulson et al., 2011; Smallegange & Coulson, 2013; Vindenes & Langangen, 2015). An alternative, to date largely unexplored methodology, is the use of systems approaches to forecast patterns at the metapopulation scale from lower-order individual variation in life history traits (Evans et al., 2013; Evans, Norris, & Benton, 2012). Because metapopulation dynamics are by definition determined by processes at lower individual hierarchy, individual-based models are highly appropriate to perform in silico experiments in systems where sound in vivo experiments are impossible to conduct at the required detail and replication. Especially when data are sparse, difficult, and expensive to collect, systems approaches enable the development of a predictive ecology, which embraces stochasticity (Travis et al., 2013).

We follow this philosophy with as specific aim to test how observed evolved variation in life-history traits affects metapopulation dynamics. We build on earlier experimental work demonstrating a profound impact of metapopulation typology on both local and metapopulation-level demography, physiology and life-history evolution in a model species, the spider mite *Tetranychus urticae* (De Roissart et al., 2016; De Roissart, Wang, & Bonte, 2015). We developed and analysed an individual-based metapopulation model closely parameterised to the experiment outlined above. The metapopulation model does not incorporate adaptive dynamics, but instead generates metapopulation dynamics in response to the empirically derived evolved individual-level life history variation. In this way, we show that the evolved trait variation especially impacts metapopulation size, but that most variation in the metapopulation dynamics is related to spatial forcing. Virtual invasion experiments demonstrated subsequently the adaptive value of evolutionary change in individuals evolving in spatiotemporally variable (classical) metapopulations, but initially not for individuals with an evolutionary background in the other metapopulation types. By including conditional stress tolerance, evolved strategies showed adaptive value in all metapopulation types.

## Methods

### A short overview of the preceding experimental evolution

In the experimental evolution (De Roissart et al., 2016, 2015), spider mites from one genetically variable source population were placed in one of the following metapopulations (1) nine patches that received every week a constant and equal amount of resources during the entire experiment (*patchy metapopulation*), (2) nine patches with the same amount of resources randomly assigned to each patch every week until the metapopulation carrying capacity was reached, hence imposing strong spatiotemporal variation in habitat availability (*classical metapopulation*), (3) six patches of which three fixed patches got every week twice the amount of resources(*mainland-island metapopulation*). As spider mites are herbivores and feeding on bean leaf discs, patch sizes were manipulated by changing the size of the leaf discs. Hence, resource availability is directly related to patch size. Patches consist of leaves of 20 cm^2^, or 40 cm^2^ for the large patches in the mainland-island metapopulations. Patches were isolated from each other by a tanglefoot© matrix preventing mites from dispersing by walking, but a wind current (2 m/s) facilitated aerial dispersal of the mites two times a week during the entire period of the experiment. Because of the passive aerial dispersal, all patches are equally isolated from each other since probabilities of immigration were not distance-dependent, only area-dependent (De Roissart et al., 2015). Metapopulation resources were renewed weekly by adding fresh bean leaves according to the treatment. No leaves were removed before complete deterioration preventing the enforcement of extinction. Dispersal in the metapopulation was thus random regarding immigration and at high cost. The metapopulation-level food availability as defined by the total availability of leaf surface over subsequent generations was identical among treatments.

An overview of the measured metapopulation metrics and evolved individual life history traits and can be found in Table 1. We found the total metapopulation size to be lowest in classical metapopulations due to the higher patch extinction rates. Variability in the local population sizes over time (alpha variability), variability in the total metapopulation size over time (gamma variability), spatial synchrony and local patch extinction rate were lowest in the mainland-island metapopulations. Alpha variability was highest in the classical metapopulations. Fecundity rates evolved towards higher values in the classical and mainland-island metapopulations than in patchy metapopulations. In classical metapopulations, longevity evolved to the lowest values, while time till maturity and male-bias among progeny increased by evolution in the mainland-island metapopulations relative to the other typologies. We additionally found an enhanced stress resistance in mites that evolved in both mainland-island and classical metapopulations relative to those evolving in patchy metapopulations.

**Table 1:**
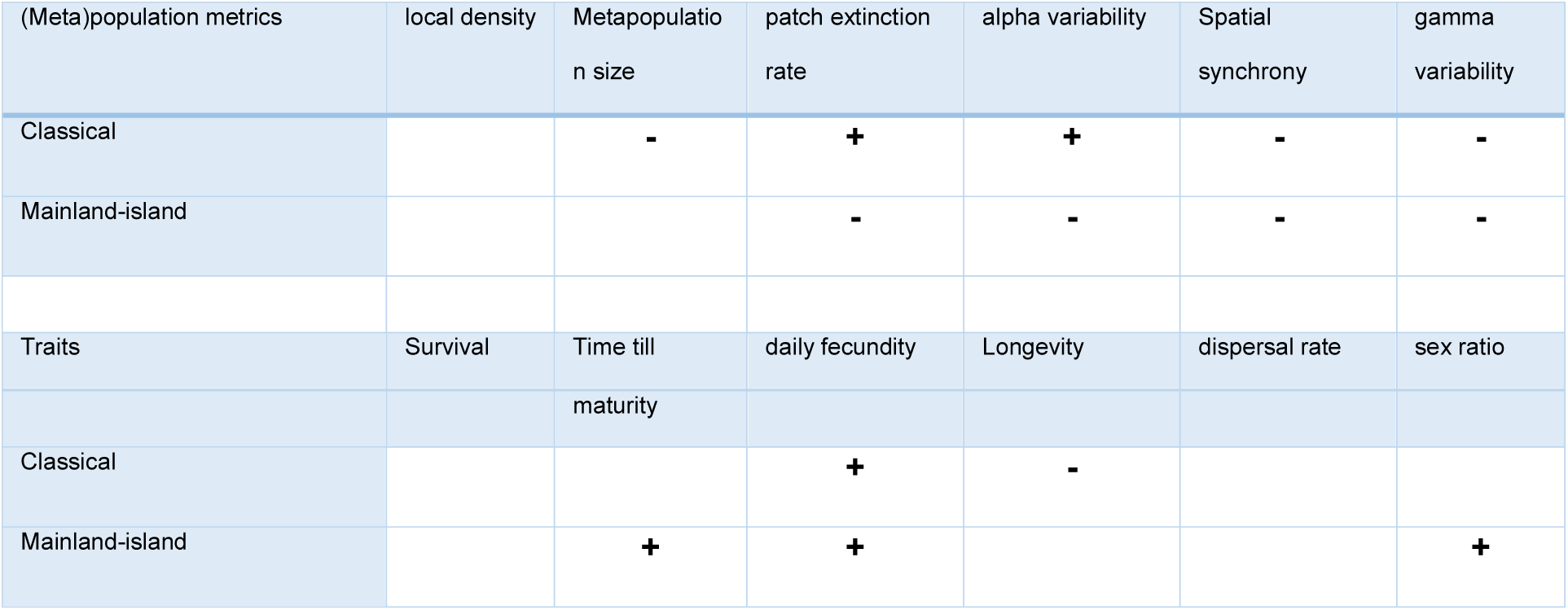
Schematic overview of the observed metapopulation dynamics and evolutionary change in life history traits in experimental classical and mainland-island metapopulations relative to patchy metapopulations (De Roissart et al., 2016). + and - indicate respectively statistically larger or smaller values of the respective (meta)population metrics relative to those measures in the patchy metapopulation. Empty cells depict represent no statistically differences in the metrics.

### The model algorithm

The model is individual-based and simulates mite development and population dynamics as in the experiments (De Roissart et al., 2016, 2015) on a daily basis for 1200 days (Fig. 1). The same spatial setting as in the experimental evolution was used as a boundary condition. The simulated individuals live according to the measured life histories and their interactions with resources availability (feeding and emigration rates). As in the experimental metapopulations, dispersal is non-directional and any virtual individual engaging in dispersal has a small and equal probability of ending up in any randomly chosen different patch.

**Figure 1:**
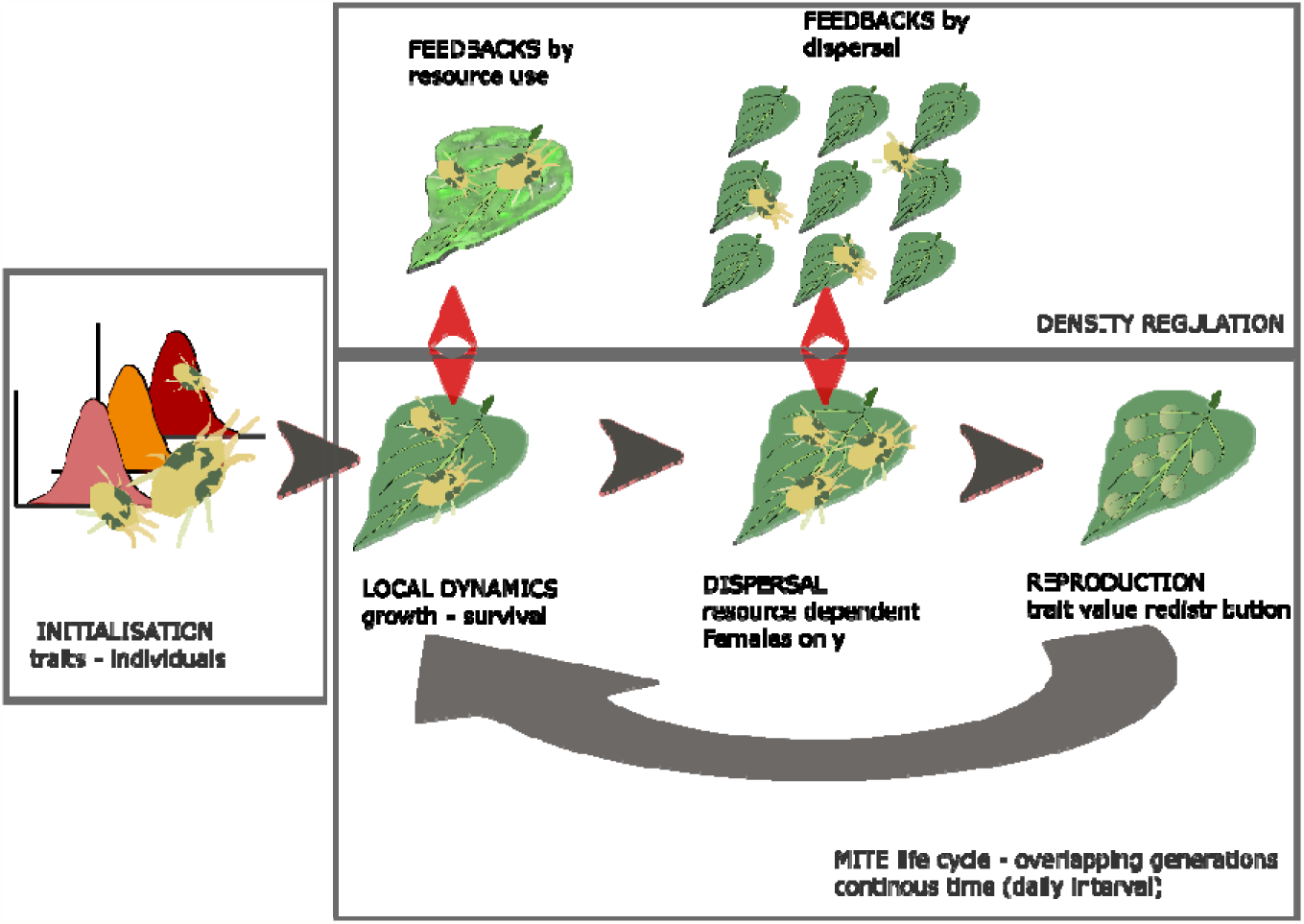
Schema of the implemented modelling routines and feedbacks according to the mite life cycles in the experimental system

The evolved traits, as determined at the end of the experimental evolution (De Roissart et al. (2016) and Table 1) were used to parameterize the model. Individuals carry alleles that determine survival rate during development, age at maturity, longevity, probability of being female (sex ratio). A total of 900 mites with trait variation as quantified by (De Roissart et al., 2016) are initialized in each metapopulation (100 individuals per patch unit). Parameterisation of feeding rates and thresholds of starvation were based on simple experiments where different stages (larvae, nymphs, adult males and females) of mites were allowed to feed on fresh bean leaves or left to survive without food. Feeding rate was assessed by quantifying the surface of bean leaves with chlorophyll removed by the different stages during one day. These experiments were separately performed with mites not originating from the experimental metapopulations and thus reflect values under standard conditions. We used approximate Bayesian methods for individual-based models (see Supplementary material 1) to demonstrate that this restriction in terms of parameter quantification did not lead to deviating metapopulation and population statistics relative to those experimentally observed (i.e., the posterior parameter ranges as determined by pattern-oriented modeming included the observed and implemented parameter values). An overview of the parameter setting can be found in Table S1 in supplementary material. All females in the metapopulation are expected to mate, so to have female offspring. To avoid selection during the simulations and thus to maintain the experimentally quantified genetic variation no genetic algorithms were implemented. Instead, new-born mites received a random trait value from the appropriate distribution. Dispersal is modelled as a global dispersal in 1-2 day old females among all patches at the experimentally derived rate and dispersal cost. As in the experimental metapopulations, dispersal is only possible every three days. Besides these experimentally derived individual measures, additional density dependency in dispersal and survival (as measured at the population level in De Roissart et al. (2015)) were implemented. Resources were refreshed as in the parallel experiments and feedbacks between population size and stage structure and resource use were thus emerging processes. As for the empirical experiments that served as base for the model, we calculated the following metrics related to demography and metapopulation dynamics (Table 1): mean local population size, mean metapopulation size, patch extinction probability, local variability in population size (alpha variability), spatial synchrony in population sizes and metapopulation-level variability of the entire metapopulation size (gamma variability). The model predicted differences in metapopulation dynamics well; the ODD-based model description can be found in Supplementary material 1.

### Virtual experiments

#### 1. Reciprocal metapopulation translocation experiment: partitioning the effects of trait variation and spatial forcing for metapopulation dynamics

Are evolved strategies to the benefit of the metapopulation? We initialized individuals characterized by trait distributions that evolved in each of the experimental metapopulation types in each of the three metapopulation types. So, 30000 simulations were run for each metapopulation scenario, 10000 with trait values that evolved in the matching metapopulation, and 2 x 10000 with trait distributions from mites evolved in the alternative metapopulation context. We prohibited evolutionary dynamics by reassigning individual trait values every generation from the empirically observed distribution rather than passing them from parents to offspring. The resulting metapopulation-level properties were calculated (see Table 1) and enabled us to:

a. quantify the importance of evolutionary background (individual variation as derived from experimental evolution) versus spatial forcing (metapopulation structure) on local and metapopulation-level demographic metrics. We partitioned variance in all metapopulation metrics by means of constrained ordination (RDA; Oksanen, Blanchet, Kindt, Legendre, & O’Hara, 2016).
b. assess to which degree metapopulations initialised with mites from the matching evolved experimental metapopulation performed better than those from the non-matching metapopulations. If evolution of individual mites is of benefit to the metapopulation, local-and metapopulation size is expected to be maximized, extinction probability, local and metapopulation-level variability and spatial synchrony to be minimised. We thus interpret metapopulation statistics as a proxy of metapopulation fitness.

#### 2. Invasion experiment

Are evolved strategies to the good of the individual? We expect evolution in the experimental metapopulations to increases individual fitness, so to be adaptive at the individual level. We therefore tested whether individuals with trait values that evolved in a specific metapopulation context (typology) were able to invade their corresponding and, hence, matching metapopulation type occupied by individuals that carry trait values that evolved in one of the two contrasting metapopulation types. The ‘matching’ mutants were consequently introduced into a fully functional metapopulations (pseudo-equilibrium reached at 600 time steps) initialised with mites from a different, and hence, non-matching evolutionary context. Similar to evolutionary invasion analyses, increases in population size of the mutant at short and longer time frames then demonstrate the adaptive value of the evolved individual strategy. A total of 90 (10/patch) of these individuals were introduced in contrasting metapopulations of ~20000 individuals to minimise stochastic extinctions after introduction. 1000 independent runs per invasion scenario were completed.

Because transcriptomic and physiological analyses in the experimental mite metapopulations indicated evolved stress resistance in mites from classical and mainland-island metapopulations (De Roissart et al., 2016), we additionally performed invasion experiments by allowing 10, 25 and 50% lower mortality rates under food shortage. This enabled us to test the prediction that such evolved stress resistance as translated in higher starvation tolerance would increase individual performance and invasibility. Invasibility was quantified as the proportion of metapopulations where the matching genotypes survived (attaining increasingly higher population sizes). High invasibility rates of the matching genotypes then indicates that the evolved life history traits are adaptive at the individual level.

## Results

### Reciprocal metapopulation translocation experiment

Evolutionary changes in life history traits explained 11.2% of the total variation in the suite of metapopulation and population demographic metrics, while 57% was explained by spatial forcing (Table 2). Especially metapopulation-level population size, and to a lesser degree local population size were influenced by the evolved trait variation (Table 2). More specifically, mites that evolved in patchy metapopulations systematically reached high local and metapopulation densities (see Fig 2).

**Table 2.**
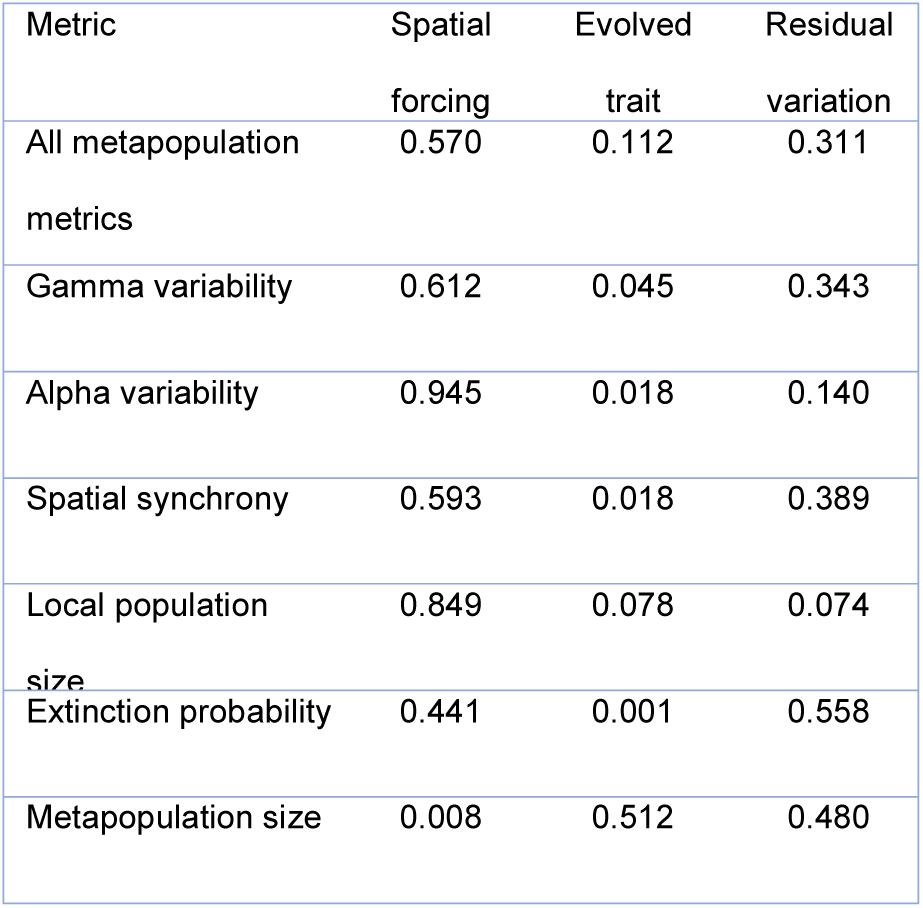
Variance partitioning of the six metapopulation metrics according to spatial forcing (metapopulation type), evolved trait variation and its covariation. The proportion of residual variation is given in the last column.

| Metric | Spatial forcing | Evolved trait | Residual variation |
| --- | --- | --- | --- |
| All metapopulation metrics | 0.570 | 0.112 | 0.311 |
| Gamma variability | 0.612 | 0.045 | 0.343 |
| Alpha variability | 0.945 | 0.018 | 0.140 |
| Spatial synchrony | 0.593 | 0.018 | 0.389 |
| Local population size | 0.849 | 0.078 | 0.074 |
| Extinction probability | 0.441 | 0.001 | 0.558 |
| Metapopulation size | 0.008 | 0.512 | 0.480 |

**Fig 2.**
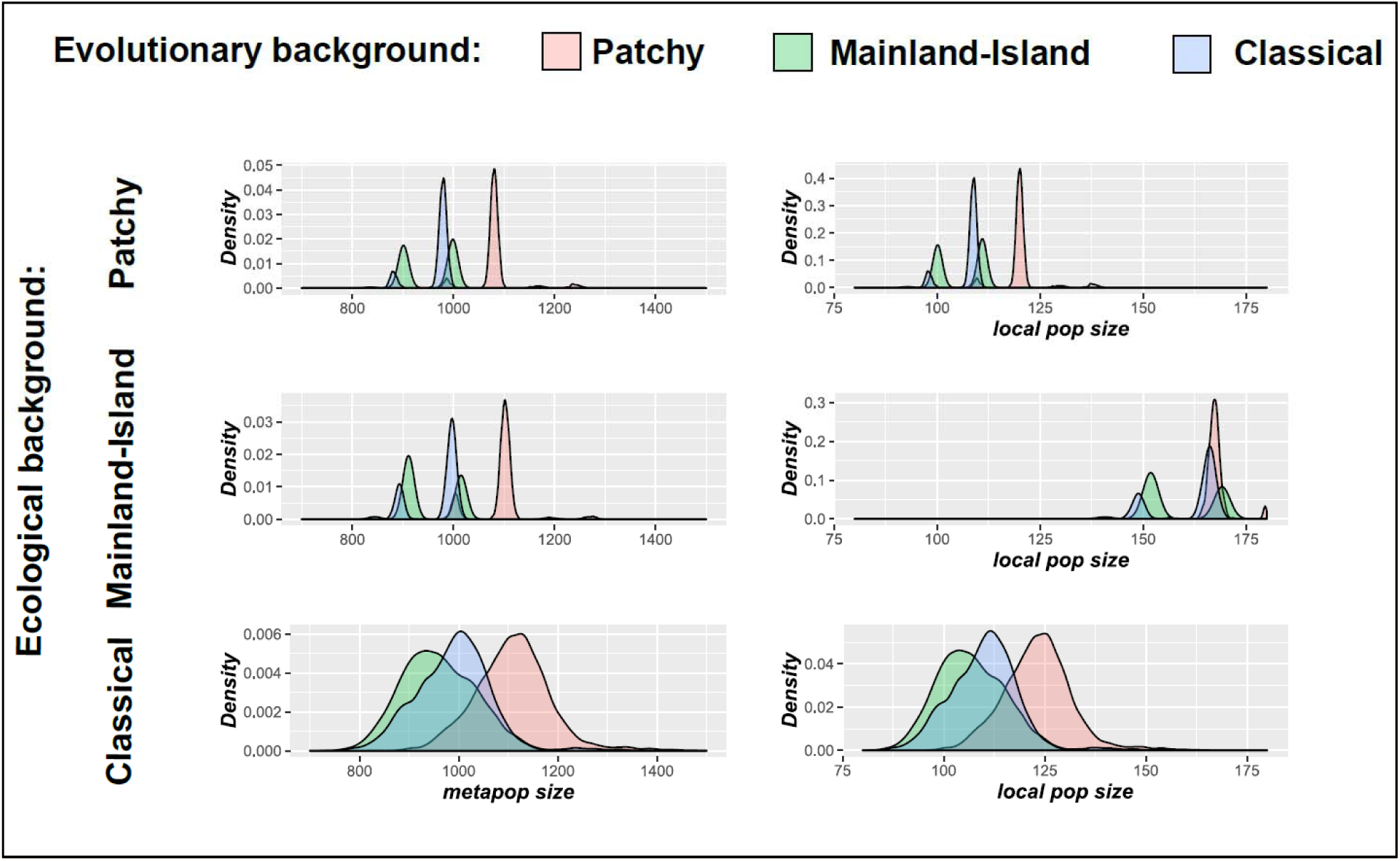
Distribution of different population sizes obtained by the virtual translocation experiments. Left panels represent the distribution of metapopulation sizes; those at the right local population sizes. Upper panels show results from virtual translocations in patchy metapopulation, Middle those from Mainland-Island metapopulations and lower ones those from classical metapopulations. Colours indicate the different evolutionary background of the introductions. The metapopulation typology (spatial forcing) is the main determinant of alpha, gamma variability and spatial synchrony. Results from virtual translocation experiments show across the metapopulation contexts highest average population and metapopulation sizes for mites with an evolutionary background from patchy populations. Population sizes are thus not maximized when genotypes match their metapopulation background

Metrics related to spatial and spatiotemporal variance in metapopulation size and local population size were predominantly impacted by spatial forcing (Table 2), with highest contributions of the spatial forcing reached for the metrics at the local level (alpha variability). Alpha and gamma variation were highest, spatial synchrony lowest in classical metapopulations (Fig 3; note the different axis-scale of alpha variability for classical metapopulations!). Spatial synchrony is highest in mainland-island metapopulations. Despite the overall low contribution of evolution to metapopulation variability statistics, gamma and alpha variability is systematically lowest for genotypes that evolved in mainland-island metapopulations, and highest for individuals with an evolutionary history from classical metapopulations.

**Fig. 3.**
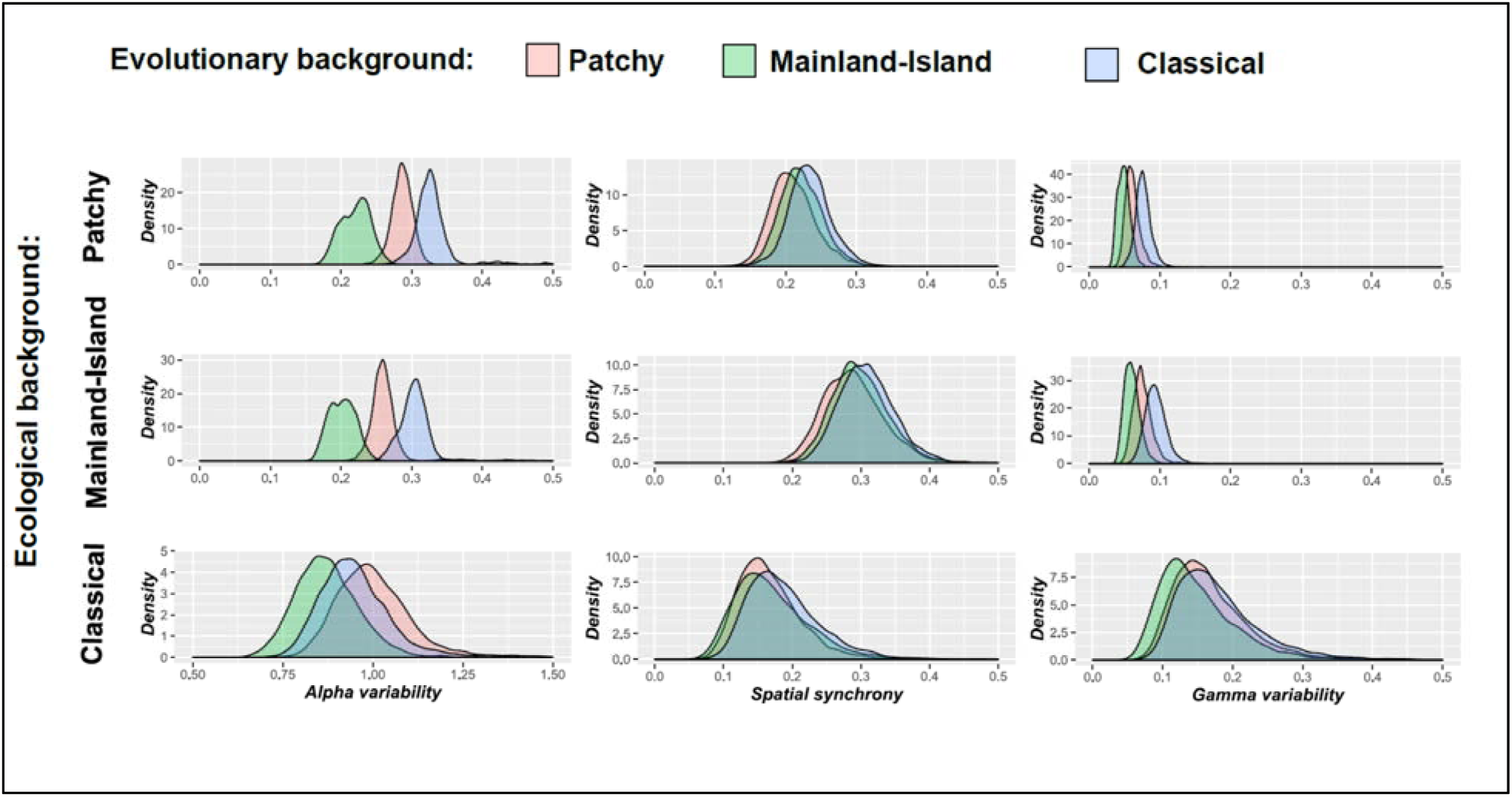
Distribution of different metapopulation variability measures obtained by the virtual translocation experiments. Panels are horizontally ordered according to the three metrics (left: alpha variability, Middle: spatial synchrony; Right: Gamma variability). Upper panels show results from virtual translocations in patchy metapopulation, Middle those from Mainland-Island metapopulations and lower ones those from classical metapopulations. Colours indicate the different evolutionary background of the introductions. The metapopulation typology (spatial forcing) is the main determinant of alpha, gamma variability and spatial synchrony. Note that ranges along X-axis is different for alpha variability in classical metapopulations relative to patch and mainland-island metapopulations.

The spatial forcing as imposed by the metapopulation typology accounted for 44% of the variation patch extinction (Table 2), while overall the contribution of evolution was marginal. This low impact of evolution is because extinctions were absent in the mainland-island and virtually absent in the patchy metapopulation (mean of 0.00012±0.00005 [0-0.0002 CI]). In the classical metapopulations, daily patch extinction risks varied between 0 and 0.028, and are overall lowest under the matching evolutionary context (mean of 0.0069±0.0055 [0-0.0244 CI]), so when populated by genotypes that evolved in classical metapopulations. Daily extinction risks were highest for individuals with an evolutionary background in the patchy (mean of 0.0084±0.0066 [0-0.0278 CI]) and the mainland-island metapopulations (mean of 0.0078±0.0066 [0-0.0256 CI]).

In conclusion, metapopulation dynamics are predominantly shaped by spatial forcing, but evolved trait variation may induce shifts in metapopulation size. We found no indications that evolution maximised population sizes or minimised local and metapopulation-level variability and spatial synchrony in the matching context. Reduced rates of extinction risk and alpha variability in the matching context for classical metapopulations indicate, however, a shift towards adaptations to the good of this metapopulation type.

### Invasion experiments

Mites that evolved in mainland-island metapopulations were least able to invade their matching metapopulation context. Genotypes that evolved in patchy or classical metapopulations had highest probability to invade their matching context, especially when the metapopulation was populated by genotypes with an evolutionary background from mainland-island metapopulations (Table 3).

**Table 3:** invasiveness of genotypes into their matching metapopulation occupied by individuals with a non-matching evolutionary history. The rates thus indicate the proportion of successful invasions of matching genotypes into their own metapopulation context populated by individuals that evolved in a different metapopulation type.

| **Metapopulation context** | **Evolutionary background of the metapopulation** |  |  |
| --- | --- | --- | --- |
|  | **Patchy** | **Mainland-island** | **Classical** |
| **Patchy** |  | 0.633 | 0.292 |
| **Mainland-island** | 0.028 |  | 0.004 |
| **Classical** | 0.233 | 0.686 |  |

As mentioned, we observed physiological stress responses in mites that evolved in the classical and mainland-island metapopulations. We therefore performed additional invasion experiments where we implemented a 10, 25 and 50% lower mortality rate under starvation in genotypes from these metapopulations. Implementation of a starvation resistance of 10% in mites from classical metapopulations, increased their invasibility in the matching context already towards 84.2% when the metapopulation was occupied with mites from mainland-island metapopulations and towards 57.3% when occupied by mites from patchy metapopulations. These invasion rates increased to more than 95% under a starvation resistance regime of 25 and 50%. The invasibility of mites that evolved in mainland-island metapopulations increased with 10, 25 and 50% lower mortality rate from 37 and 32 over82 and 52 towards 100 and 87.5% when they were respectively invading their matching context occupied by mites with an evolutionary history from patchy and classical metapopulations. Implementing starvation resistance thus also increased the adaptive nature of the life history evolution, especially in mites that evolved in mainland-island metapopulations.

## Discussion

Changes in the spatial structure of populations have a strong impact on demography and genetics. Theory predicts that a coupling between ecological and evolutionary dynamics may affect functioning and performance of the metapopulation (Ezard, Côté, & Pelletier, 2009; Hanski & Mononen, 2011; Kinnison & Hairston, 2007). Understanding to which degree metapopulation dynamics are affected by evolutionary dynamics is essential for conservation, as conflicts between individual and metapopulation goods may both increase or decrease metapopulation performance in terms of persistence and size, and eventually lead to evolutionary rescue or suicide (Delgado et al., 2011). Because local populations are connected by dispersal in metapopulations, dispersal is expected to be a central life history trait for these eco-evolutionary dynamics (Bonte & Dahirel, 2017). Many theoretical, but also experimental studies using bacterial and protist populations confirm this view (Bell & Gonzalez, 2011; Fronhofer & Altermatt, 2015). In reality, evolutionary dynamics are not restricted to dispersal evolution, but comprise multivariate selection on a wide array of traits, as earlier reported for our experimental system (De Roissart et al., 2016). By building further on this experimental work using an arthropod model, we here show that about 40% of the metapopulation dynamics, as measured by metrics related to population size means and variances, are determined by the spatial setting *per se*. From these performance metrics, average local population sizes are about 8% determined by evolutionary changes in life history traits, while the variation in total metapopulation sizes is up to 50% explained by evolved life history changes. Overall, our estimates of the contribution of evolution to population sizes are in line with those estimated for phenotypic change in freshwater crustaceans and rotifers and for community and ecosystem response following guppy evolutionary change after predator introduction in Trinidadian streams (Ellner, Geber, & Hairston, 2011).

Evolutionary changes in life history explained between 8% and 50% of all variation in population sizes among the three experimental metapopulation types. Although the magnitude of this impact is the result from our chosen spatial settings, it nevertheless demonstrates the potentially strong impact of evolution on the eventual reached population sizes in metapopulations. As metapopulation size was largely constrained by the experimental setup and deliberately fixed for all treatments, it is not surprising that the contribution of spatial forcing is low. It remains nevertheless surprising that the effectively reached population sizes depends on the evolutionary history. While a profound theory is needed to further explore the exact causes of these eco-evolutionary dynamics [Govaert et al. this issue], theory is hinting at the underlying mechanisms (Zhang et al. 2017). The evolution of life histories creates strong heterogeneity in the local environmental conditions due to feedbacks in local demography and dispersal (Bonte et al. 2018). According to theory, such feedbacks may pull the metapopulation-level carrying capacity beyond the one reached by summing the local population sizes if they were isolated from each other (De Angelis, Ni & Zhang 2016; Holt 1985). Feedbacks from life history on local-and metapopulation demography and on spatial variation in local environmental conditions (resource availability through density changes) are hence sufficient to impose changes in metapopulation-level carrying capacities. The finding that virtual mites that evolved in patchy metapopulations reached highest local and metapopulation sizes overall does demonstrate the validity of this eco-evolutionary process. Under this spatial evolution regime, evolved strategies indeed lead to enhanced connectivity and spatial heterogeneity following the evolution of respectively increased dispersal and growth rates.

The observed life history evolution in mainland-island and classical metapopulations did consequently not optimise population sizes because they were not largest in their matching context. Because demographic variability was predominantly determined by spatial forcing, variability and spatial synchrony were neither found to be lowest under matching scenarios. Local and metapopulation-level variability was overall lowest for genotypes that evolved within mainland-island metapopulations. While local extinction rates were predominantly determined by spatial forcing and only relevant in classical metapopulations, a reduced rate was found for matching genotypes. Life-history evolution thus improved metapopulation dynamics that improve persistence to some degree only in our classical experimental metapopulations.

The predicted local population and metapopulation sizes under genotype-metapopulation matching accord with those obtained from empirical work. Although earlier experiments were only replicated three times, it is interesting to note that similar bimodal patterns in local population and metapopulation sizes were found in the experiments. These measures are time snapshots at the end of the experiment. The bimodal patterns result from temporal fluctuation with crowded patches dominated by long-lived adult females alternating with freshly colonised patches dominated by juveniles. Our model systematically predicted highest metapopulation sizes for genotypes from patchy metapopulations, independent of the metapopulation structure. Differences in population and metapopulation sizes across treatments are in line with those empirically derived (De Roissart et al., 2015) but differences between patchy metapopulations on the one hand, and mainland-island and classical metapopulations on the other hand are larger than observed.

We only modelled variation in the experimentally derived life-history traits (De Roissart et al., 2016). These were all quantified under optimal conditions. Our invasion models suggest that the incorporation of conditional mortality, expressed as starvation resistance for genotypes originating from classical and mainland-island metapopulations turned evolutionary life-history changes adaptive. Decreasing starvation mortality in the invasion experiments, gradually lowered differences in predicted average population sizes relative to those from patchy metapopulations and decreased variances for metapopulations populated by these genotypes, but not to such a degree that overall patterns changed towards optimised metapopulation functioning under matching contexts. Time and other practical constraints make it difficult for empiricists to measure life-history traits under different density conditions. An increased tolerance to starvation is, however, only one of the possible evolved conditional strategies related to a general stress resistance. Such a general stress resistance strategy was revealed by detailed transcriptomic and physiological analyses. Unfortunately, we did not obtain the results from these molecular analyses whilst the experiments were running, so we were unable to perform additional starvation or other stress resistance tests, except for the performance tests on a subordinate host plant (tomato), which in line with the molecular analyses showed a better performance for mites that evolved in the classical and mainland-island metapopulations. Other density-and condition-dependent strategies may equally generate feedback on population sizes. Changes in feeding efficiency as implemented into functional responses in a resource-consumer model for instance predicted elevated densities of protists in a range-expansion experiment (Fronhofer & Altermatt, 2015).

While the link between life-history evolution and metapopulation functioning is not straightforward, selection should maximise fitness and thus enable genotypes to invade their matching environment occupied by genotypes that have not evolved locally. Our invasion experiments confirmed this principle for individuals with an evolutionary background in patchy metapopulations. Virtual mites with traits from mainland-island and classical metapopulations were, however, not able to invade their matching metapopulation context, especially when these were occupied by genotypes from the patchy metapopulation. This apparent maladaptive evolution could be countered by translating the observed signals of stress resistance into reduced mortality rates under starvation. 10-25% lower mortality rates under starvation led to successful invasions. Our study thus indicates that even small conditional changes in particular life history traits in spider mites, which were in line with insights from–omic approaches (De Roissart et al., 2016), were key to optimality under matching demographic conditions. This also implies that our estimate of the contribution of evolutionary change to metapopulation dynamics is likely a lower and conservative boundary. The overall level of explained variance by the spatial setting and evolution was, however, little affected when stress resistance was implemented (see Supplementary material 2).

We show that in our spider mite experimental metapopulations, both temporal variation in local and metapopulation size, as well as spatial synchrony are predominantly determined by spatial forcing as imposed by the spatial configuration of the metapopulation. Local and especially metapopulation-level average population sizes are in contrast more determined by evolutionary dynamics. This finding implies that changes in spatial networks have the ability to affect spatiotemporal fluctuations in population sizes, thereby potentially compromising metapopulation viability, for instance by increasing population synchrony (Walter et al., 2017; Wang et al., 2015). Evolutionary dynamics may in contrast increase average population sizes thereby rescuing them from demographic and environmental stochasticity. These theoretical insights should in first instance be regarded as a proof of concept demonstrating that changes in spatial configuration in metapopulations may affect different properties of the metapopulation dynamics more than evolution resulting from these changes in spatial structure, and that the contribution of evolution to metapopulation dynamics can be substantial.

Obviously, the presented insights could not be obtained from adaptive dynamic modelling. Such approaches typically focus on the optimisation of single, or few traits at maximum [Govaert et al.; this special issue] while neglecting the complex architecture of the multiple life history traits that eventually determine the species’ life history. The genotypic variance/covariance matrix is unknown in many, not to say all species. Feedbacks in eco-evolutionary ecology are complex and often involving non-linear dynamics and conditional processes. While analytical and traceable eco-evolutionary models are essential to further develop a sound theoretical framework to understand the putative importance of evolutionary dynamics on ecological processes [Govaert et al.; this special issue], we advocate the development of systems modelling as a mature discipline in ecology that encapsulates both the complexity and the multivariate nature of life history evolution and the importance of individual-level variation for higher-level ecological processes. Such a development per definition crosses the boundaries of theoretical biology, empirical ecology and computational sciences and should speed-up the transition of the field of eco-evolutionary dynamics towards a predictive science.

At the scale of the experimental setup, our results point to the fact that changes in the arrangement of host plants, for instance in horticulture, may strongly affect densities and damage thresholds for pest species, both directly through spatial forcing and the resulting genetic changes in life history traits and physiology. In a broader perspective, we show that local population sizes of species inhabiting metapopulations may benefit from metapopulation-level evolutionary changes in traits that are not traditionally linked to metapopulation persistence.

## Acknowledgement

We are grateful to Annelies De Roissart and Katrien Van Petegem for providing the life history data needed to parameterise the model. The project was funded by FWO (G.018017N); QB is a holder of FWO doctoral fellowship. D.B. was supported by the FWO research network EVENET (grant W0.003.16N). We finally would like to thank reviewers involved in PCI Evolutionary Biology, the organisers of this special issue and all members of the TEREC journal club for reading and commenting earlier versions of our manuscript.

## Author Contributions

DB conceived the ideas, designed the methodology and led the writing of the manuscript; QB coded the model. DB & QB contributed critically to the drafts and gave final approval for publication.

## Supplementary material 1: ODD of the model

The model is individual-based, object oriented and coded in Python 3.5. Code, including parameterization for the three metapopulation types, is provided on Github https://github.ugent.be/pages/dbonte/eco_evo-metapop/

The model description follows the ODD (Overview, Design concepts, Details) protocol for describing individual-and agent-based models (Grimm et al. 2006, 2010).

### 1. Purpose

The purpose of the model is to simulate the population dynamics of spider mites in three types of metapopulations (De Roissart et al. 2015). The model simulates spider mite demography in these metapopulations based on empirically retrieved life history parameters (De Roissart et al. 2016) at the end of the experimental evolution. Population dynamics as simulated by the model are thus hierarchically dependent on the <u>implemented spatial forcing through differences in metapopulation typology</u> (the spatiotemporal variation in patch sizes as an externally spatial forcing variable) and the <u>evolved trait variation</u> as observed in experiments (including stochastic variation). By following a reciprocal virtual experiment, the model allows to test how much of the variation in metapopulation demography can be attributed to the spatial forcing or the evolved trait variation. It additionally allows a rigorous test on whether trait evolution maximizes <u>metapopulation performance</u> in terms of population sizes and variance, under the matching context, i.e. when virtual metapopulations are populated by individuals with traits that evolved in this spatial context. The model is also used to test the hypothesis that individuals are able to <u>invade</u> their matching metapopulations occupied with genotypes that evolved in one of the non-matching metapopulations.

### 2. Entities, state variables, and scales

The model contains the following **entities**: *individual mites*, *local patch* in the metapopulation and *metapopulation typology*, each with their specific attributes and states:

The *metapopulation* has three states: either it consists of (a) nine, equally sized patches that are constant in size in time (homogeneous metapopulation; or ***patchy metapopulation***; abbreviated as HOM), (b) six patches of which are three of double size as the other and constant in size in time (Spatial heterogeneous metapopulation or ***mainland-island metapopulation***; abbreviated as SPA) or (c) a set of patches that are of variable size in space and time but in total reaching the same metapopulation capacity as (a,b). Here, patches can thus being either of double size or absent (spatiotemporal heterogeneous metapopulations, or ***classical metapopulation***; abbreviated as TEM). Total metapopulation carrying capacity, as determined by the summed size and hence resource availability of the local patches is always the same. In the experimental setup, local patches were bean leaves of standardized size, the surface determines the availability of resources (cells with chlorophyll) for the mites. The model is run for 1200 days.

The *local population* has as states the total number of resources that us weekly assigned to the specific patch.

*Individuals* are characterized by the following attributes that are

1. fixed during the simulation at the individual level: sex at maturity, maximal age-dependent resource use (plant cells of unit 1mm^2^/day), age-dependent daily mortality rate, daily fecundity (eggs/day), time till maturity (days) and resource-dependent emigration strategy. All these life history traits are sampled from predefined Gaussian distributions based on experiments.
2. continuously updates (daily basis) throughout the simulations: age, condition as determined by food uptake and the local patch it inhabits.

The model’s **spatial scales** are thus determined by the number and the size of the local patches, and are the same among the three metapopulation types. The size of the patches represent the size of the leaf surface on which mites live (mm^2^). Mite demography as determined by feeding, aging, egg-laying, dying and dispersal is modelled at a daily basis.

### 3. Process overview and scheduling

Individual mites live their life in the metapopulation on a daily basis. At every time step, individuals age (from egg to adult) and age is counted as number of days since eggs were deposited by the parents. Depending on the individually assigned traits at birth, individuals will continue to live, take-up the maximal number (according to their age and sex; males do not feed anymore) of the available resources, disperse according to their age and the local quantity of available resources and lay eggs according to their age (adult females) and availability of resources, or die. These processes are sequentially run in randomly selected mites, so the number of resources is continuously updated. Emigration is as for the accompanying experiment, only possible at three-days intervals (simulating temporal variation in external suitable wind conditions for aerial dispersal). The number of mites per patch is recorded after the execution of the daily routines for all individuals.

The number of resources in each patch is weekly refreshed according to the predefined metapopulation typology.

### 4. Design concepts

#### Basic principles

The model simulates the local and metapopulation-level population dynamics of spider mites in predefined metapopulations. Individual lives are daily updated, and collective feedbacks are emerging on the available resources. Density regulation following competition emerges from mortality by starvation, density regulation by means of dispersal is equally resource dependent and affected by immigration costs as settlement rates are proportional to the available habitat and low (5%).

#### Emergence

Populations are building-up in the set of connected populations. Both spatial and temporal changes in population sizes and age structure are emerging.

#### Adaptation

The model does not contain any adaptive dynamics. All incorporated individual variation stems from observed variation in the parallel *in vivo* experiments. This implies that at birth, parameters are sampled from the same statistical distribution. Trait values are hence not inherited.

#### Objectives

As no adaptive traits are explicitly modelled, individual’s success is not modelled to meet certain objectives

#### Learning

No learning algorithms were implemented

#### Prediction

Individuals do not engage in adaptive decision-making

#### Sensing

Individuals condition their departure decisions based on the observed amount of patch-level resources. individuals die faster when local resources become limited according to age-dependent resource thresholds

#### Interaction

Individuals interactions indirect via competition for a resources.

#### Stochasticity

The life history traits associated with individuals is sampled at birth from empirically defined Gaussian distributions. Further, all chance events (survival, emigration, sex ratio) are run for all individuals at the relevant temporal scales (daily basis for survival, sex-ratio at birth, dispersal during the early adult phase). Because of feedbacks with resource availability, stochastic population regulation is emerging through imposed starvation. In the classical metapopulation (TEM), resources are weekly and randomly associated to patches in the metapopulation.

#### Collectives

No collective behaviours are modelled.

#### Observation

At the end of the simulation, metapopulation statistics are calculated: mean local and metapopulation size, temporal variability of local subpopulations (alpha variability) and the metapopulation (gamma variability), synchrony of local subpopulation size changes. Metrics are calculated following Wang et al. (2014). These measures of metapopulation performance are expected to vary according to the implemented individual trait variation.

### 5. Initialization

The metapopulations are initialized as a network of either six (SPA) or nine patches (HOM,TEM) with the maximum level of resources (expressed as mm^2^ chlorophyll, or leaf surface). In the metapopulations with nine patches, all patches have as starting value 5000 for the resource level. In the SPA, three patches have a value of 10000. So metapopulation-level resource availability is equal in all simulations. The metapopulation is initialized with 100 individual mites per patch-unit. The trait values for each individual are sampled from distributions as observed in the experiments (De Roissart et al. 2016). Each individual is randomly assigned to an age (between 3 and 11 days).

### 6. Input data

Except for the mentioned individual trait parameterization, the model does not contain input from external sources.

### 7. Submodels

#### Local dynamics

##### Mite daily life: the LIVE procedure

Individual mites have the following attributes: the metapopulation to which it belongs, its local patch number (location in the metapopulation), its age, sex, resource level and metapopulation-specific (so evolved) life history traits according to those observed in the experimental evolution experiment: mean sex ratio, mean fecundity, mean dispersal intercept relative to resource level, mean juvenile mortality, mean longevity as adult, mean juvenile mortality and time till maturity. Age-dependent resource use (maximum uptake of resources) is determined from parallel experiments and assigned to individuals independent from their genetic background. All values are sampled from 95%-truncated Gaussian distributions to ensure values within the range of observation and avoiding negative values.

Overview of the empirically determined metapopulation-independent traits in parallel lab assays (not published).

- mean resource use as adult female: 4.12 (1.14 SD) mm^2^/day
- mean resource use as nymph 0.616 (0.20 SD) mm^2^/day
- mean resource use as larvae: 0.31 (0.11 SD) mm^2^/day

Aerial dispersal is modeled as a resource dependent strategy, only occurring when the leaves are not depleted of resources (set to 1500 mm^2^). The slope is fixed for all individuals independent of their genetic background (0.122). We found the aerial dispersal propensity to increase with resource availability at the population-level (see De Roissart et al. 2015), such that the individual probability to emigrate equals InterceptDispersal+0.122*log(resources available). We used this population-level quantification rather than results from behavioural experiments under optimal conditions within limited temporal windows (e.g. De Roissart et al. 2016).

Other empirically determined life history parameters for the individuals that are specific to their evolutionary history

**Table S1.** Mean trait values and standard deviations

| Evolutionary history (metapopulation) | Daily fecundity | Sex ratio | Dispersal intercept | Juvenile mortality | Time till male maturity | Time till female maturity | Adult survival rate |
| --- | --- | --- | --- | --- | --- | --- | --- |

Table S1. Mean trait values and standard deviations
| type of origin) | (eggs/day) | (mm/total) | (/day) | (/day) | (days) | (days) | (/day) |
| --- | --- | --- | --- | --- | --- | --- | --- |
| HOM | 4.32<br>(1.48) | 0.26<br>(0.04) | 0.7 (0.5) | 0.123<br>(0.003) | 7.94<br>(0.43) | 8.25<br>(0.46) | 0.1<br>(0.003) |
| SPA | 5.41<br>(1.48) | 0.34<br>(0.04) | 0.58<br>(0.5) | 0.156<br>(0.003) | 8.21<br>(0.43) | 8.58<br>(0.46) | 0.11<br>(0.003) |
| TEM | 5.59<br>(1.48) | 0.28<br>(0.04) | 0.7 (0.5) | 0.123<br>(0.003) | 7.98<br>(0.43) | 8.31<br>(0.46) | 0.14<br>(0.003) |

Individuals have different mortality rates according to the availability of resources and age. As for dispersal, a patch-resource availability threshold is used (the level from which dehydration or wilting of the leaf is inducing starvation). When the threshold is passed, individuals have a higher mortality rate.

- Eggs die at probability of 0.9 when no resources are available
- Larvae aged <=2 days die at probability of 0.9 when resources on the leave are lower than the threshold (dehydrated leaf)
- Nymphs (>2 and not yet mature) have a daily mortality rate of 0.9 when resources < threshold; otherwise according to the juvenile mortality rate (see table S1)
- Adults on leaves with resources < threshold have a daily mortality rate of 0.32, otherwise rates according to the adult survival rate (see table S1)

In order to test to which degree increased stress resistance, as indicated by transcriptomic and physiological analyses (De Roissart et al. 2016) affected population and invasion dynamics, mortality rates were decreased with 10, 25% and 50% under starvation.

Since males do not feed as adults, they were omitted from the population dynamics when maturity was reached.

Only mature 1-2 days old females disperse aerially (De Roissart et al. 2013;Van Petegem et al. 2015, 2016). As in the experimental metapopulations, suitable environmental conditions for dispersal (wind) were only imposed every three days. Individuals disperse randomly to any other patch with a dispersal mortality of 95% (chances of ending up in the hostile matrix)

##### Mite reproduction: the REPRODUCTION procedure

Since males mate with multiple females, we did not assume any mate limitation and assumed all females to be fertilized, thereby producing offspring at a rate according to their evolved daily fecundity with a sex ratio as equally observed in the experiments (see table S1). Egg production only started two days after reaching maturity.

##### Refreshing of resources: the PATCHRESOURCES procedure

Resources are added to the patches at a weekly basis. For the HOM, each patch received 5000 units resources; for the SPA, three patches receive weekly 5000, and another three receive weekly 10000 units of resources; patches in the TEM have an equal chance to get 10000,5000 or 0 units till the metapopulation-level carrying capacity of 9*5000 resource units is reached.

##### Invasion of different genotypes: the INTRODUCTION procedure

At time 600, so when the metapopulation dynamics are equilibrated, 10 mites of a different evolutionary history of age [3-11] (so that evolved in a different metapopulation) are introduced to test their invasion capacity in terms of persistence and population growth within the metapopulation type (SPA, HOM or TEM) and the initiated genetic constitution of the individuals (metapopulation type in which they evolved).

#### life cycle and population dynamics

The life cycle of the mites is coded as close as possible to realistic mites inhabiting the experimental metapopulations. Individuals mature according to experimentally derived growth rates, and each of the stages is characterised by experimentally quantified feeding rates, expressed as mm^2^/leaf chlorophyll consumption per day. We did not implement evolved variation in feeding rates to generate evolved reaction norms. This decision was based on the fact that measurement errors are likely larger than observed variation due to genetic background (we did not find any signature of variation in feeding rates among a selection of genotypes). Individuals compete randomly for food and are assumed to forage on the entire leaf. These dynamics create a lottery competition for resources, and feedback on resource availability within the patch. When resources are below a predefined threshold level, individuals have a predefined probability to die. Mortality and feeding rates are age-dependent. The following life histories are specific to the evolutionary background and determined by a single allele: DAILY FECUNDITY, SEX RATIO, DISPERSAL PROPENSITY [intercept of the reaction norm with starvation], BASELINE JUVENILE and ADULT DAILY MORTALITY RATE, DEVELOPMENTAL SPEED for males and females and ADULT LONGEVITY. Reproduction is only possible for >2 days old adult females when sufficient resources [set to 20% of a single leaf] are available; sex of the offspring is assigned according to a sex ratio allele. We assigned the empirically derived mean trait values, allowed them to be conditional to resource state (survival) and implemented genetic variation by sampling the trait values from the appropriate Gaussian distribution. Individual variation is thus generated by both ontogenetic (innate) and developmental plasticity (age, nutritional condition)

#### Metapopulation dynamics

Only young 1-2 day-adult females disperse when sufficient resources are available (RESOURCE THRESHOLD; starved females do not invest in aerial dispersal and during two days a week, according to the implemented experimental setup (see De Roissart et al. 2015). Dispersal is conditional with probability following the empirical derived function DISPERSAL.ALLELE + 0.122 * log(RESOURCE AVAILABILITY). The reaction norm is independent of the evolutionary background.

Dispersing individuals experience immigration costs as a 95% mortality cost. If successful, dispersing females immigrate in a randomly chosen patch (see De Roissart et al. 2015).

Resources are refreshed every 7 days according to the implemented metapopulation structure. Metapopulation dynamics thus depend on the trait initialisation and spatial setting. This allowed us to monitor metapopulation dynamics in a full factorial design metapopulation TYPE OF ORIGIN (evolutionary background) * METAPOPULATION TYPE (arena of population dynamics). Three matching and six mismatching scenarios are consequently modelled. For the invasion experiment, a small number (10 individuals/patch unit; 90 individuals/metapopulation) of matching genotypes are introduced after 600 time steps in a mismatched scenario to quantify invasion probability as a measure of fitness (invasion criterion approach).

#### Metrics

We measured local and metapopulation size at the end of each simulation. Temporal variability at both local population (alpha-variability) and metapopulation scales (gamma-variability) was calculated following (Wang & Loreau 2014). Alpha-variability is calculated as the square of the weighted average of the coefficient of variation (CV) across local populations; gamma-variability, as the square of the CV of the metapopulations. Spatial synchrony is quantified as a metapopulation-wide measure of population synchrony and equals the ratio of metapopulation gamma-variability to local alpha-variability. The patch extinction probability was measured for each run by dividing the number of extinction event by the total number of patches screened over all time steps.

#### Model parameterization and parameter sensitivity

Because life traits were either measured at the individual level under optimal conditions (single mites on fresh resources), from population-level dynamics (density dependency) or from single experiments neglecting both density dependency and putative variation among the experimental populations (feeding rates) we applied Approximate Bayesian Computation for individual based models (ABC4IBM; Van Der Vaart et al. 2015; 2016) to determine to which degree these empirically derived parameters deviated from those posteriori expected based on pattern oriented modelling using flat and broad priors centred around the observed ones.

A total of 100000 simulations were run with randomly selected trait values *x*_*t*_ sampled from a uniform prior distribution 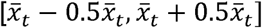 with 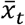 the mean observed trait value from that specific experimental setting (De Roissart et al. 2016). The resulting emerging metapopulation performance metrics (see Table 1) were calculated for each of these simulations. The deviance from observed values was calculated after z-transforming all statistics using the prior mean. Deviances of metapopulation pattern parameter D(p)were summed per simulations and provided the Goodness of Fit (GoF) for the simulation initialized with prior mean life traits

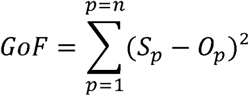

From the 100000 calculates GoF /experimental scenario, the 100 with lowest deviance were chosen, verified whether they matches the observed metapopulation patterns for each of the specific metapopulation treatment in De Roissart et al. 2015), after which the posterior distributions of the matching trait distributions were calculated median and 95CI. All posteriori distributions included the experimentally determined prior distribution, so we considered the model as robust for further use in the virtual experiments (see Table S2).

**Table S2.** Mean observed and posterior distributions of the life history parameter as determined from best matching model in terms of metapopulation dynamics as selected by Approximate Bayesian Computation.

| <u>Measured at the individual level</u> | <u>Classical</u> |  | <u>Mainland-island</u> |  | <u>patchy</u> |  |
| --- | --- | --- | --- | --- | --- | --- |
|  | <u>Mean</u> | <u>posterior</u> | <u>Mean</u> | <u>posterior</u> | <u>Mean</u> | <u>posterior</u> |
| Female age at maturity (from egg to adult) (days) | 8.31 | 6.2-10.8 | 8.58 | 6.2-10.8 | 8.25 | 6.2-10.8 |
| Male age at maturity (from egg to adult) (days) | 7.98 | 6-10.5 | 8.21 | 6.1-10.5 | 7.98 | 6.1-10.8 |
| Sex ratio (mm/tot) | 0.28 | 0.21-0.36 | 0.34 | 0.16-0.5 | 0.26 | 0.16-0.41 |
| Juvenile baseline mortality (/day) | 0.124 | 0.06-0.17 | 0.156 | 0.08-0.21 | 0.124 | 0.07-0.18 |
| Adult baseline survival (/days) | 0.10 | 0.08-0.21 | 0.11 | 0.08-0.23 | 0.14 | 0.13-0.23 |
| Dispersal propensity (intercept of the density dependent reaction norm) | 0.7 | 0.35-1.0 | 0.58 | 0.31-0.82 | 0.7 | 0.35-1.0 |
| Daily fecundity (eggs) | 5.59 | 3.3-7.8 | 5.41 | 3.2-7.5 | 4.31 | 2.9-8.1 |
| <u>Measured in independent experiments at the population level</u> |  |  |  |  |  |  |
| Density dependency of dispersal (slope) | 0.122 [0.07-0.17] |  |  |  |  |  |

## Supplementary material 2: Sensitivity of variance partitioning to unmeasured stress resistance

As for the variance partitioning of the model using the empirically quantified life history data, we additionally implemented the same procedure on models where mites from the mainland-island and the classical metapopulations received a starvation mortality of −50%, −25% and −10%. Instead of running sensitivity analyses for all traits separately, we partitioned variances for the variability statistics by jointly analysing alpha, gamma variability and spatial synchrony. Variance partitioning of the local and metapopulation size and extinction rate metrics is summarised as population size statistics.

The results are summarised below:

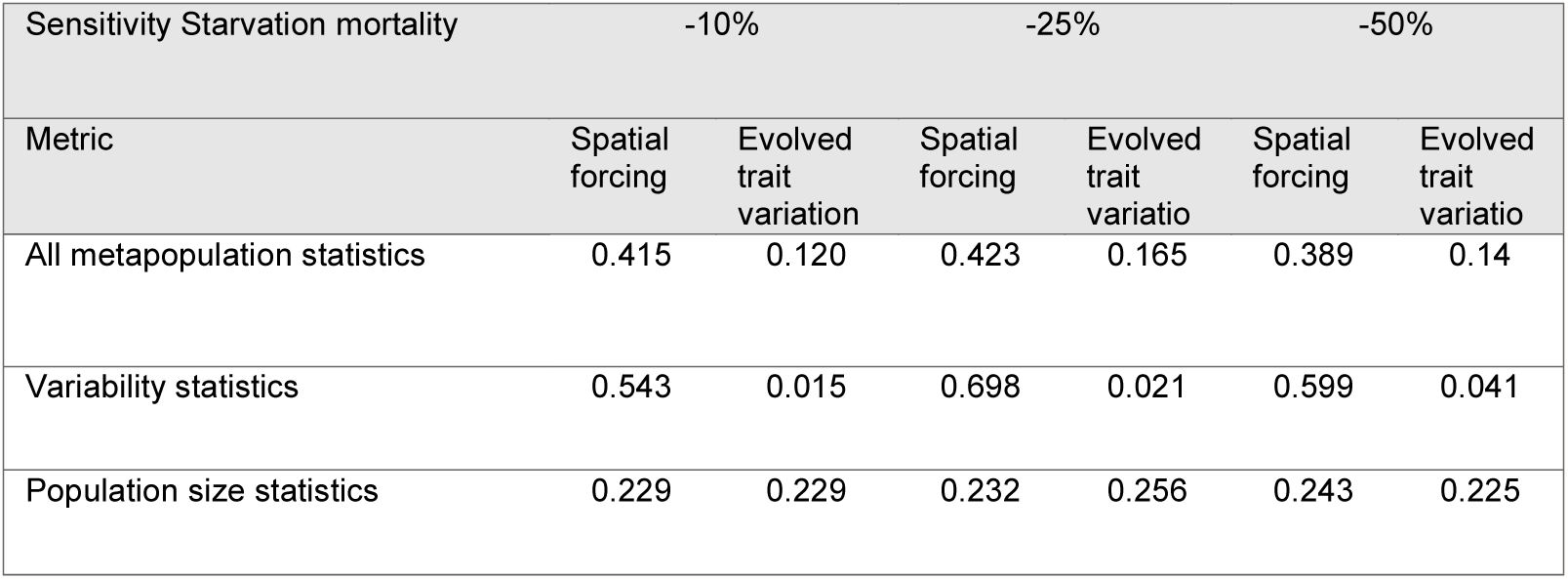

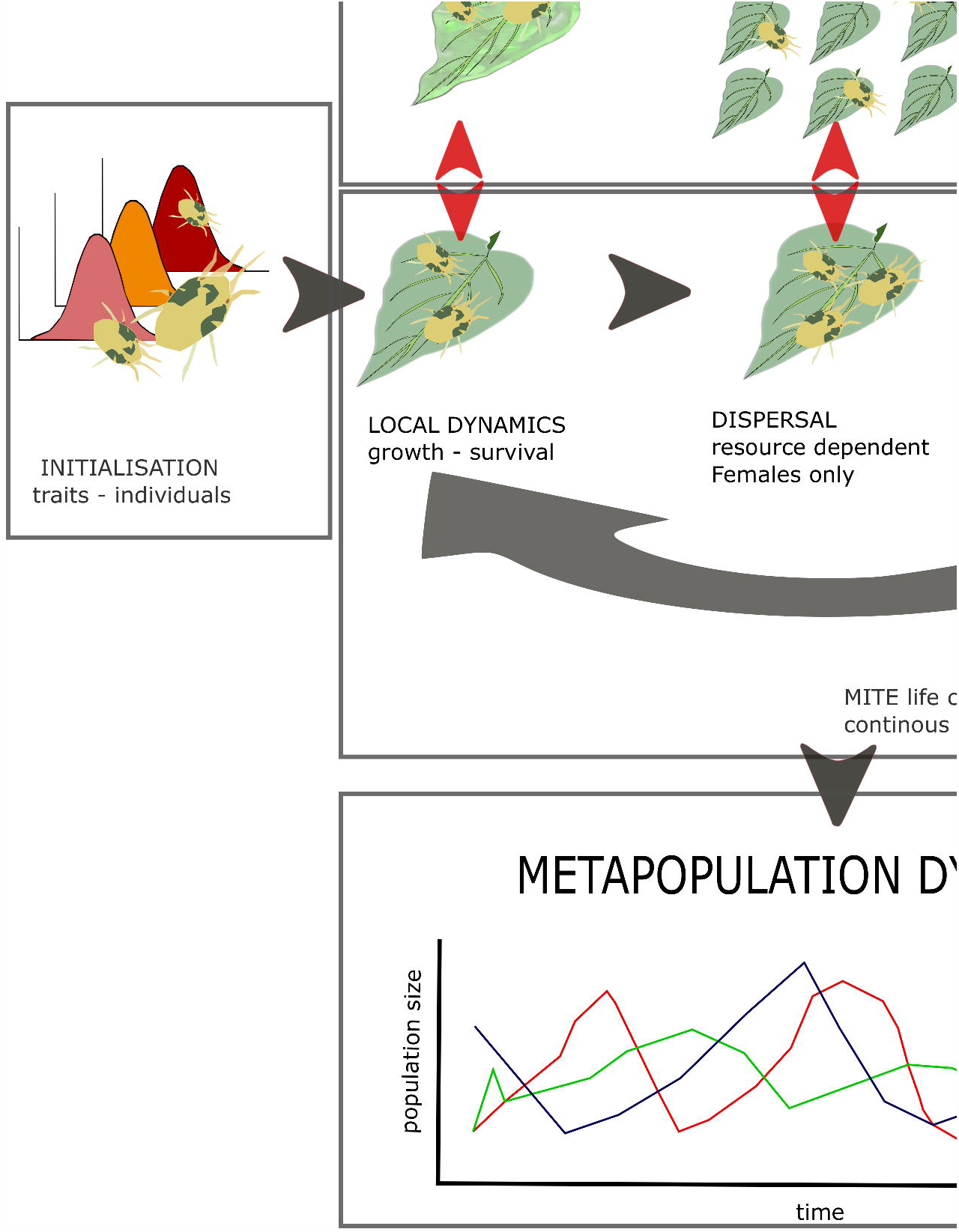

